# Diffusion as a ruler: modeling kinesin diffusion as a length sensor for intraflagellar transport

**DOI:** 10.1101/156760

**Authors:** Nathan L. Hendel, Matt Thomson, Wallace F. Marshall

## Abstract

An important question in cell biology is whether cells are able to measure size, either whole cell size or organelle size. Perhaps cells have an internal chemical representation of size that can be used to precisely regulate growth, or perhaps size is just an accident that emerges due to constraint of nutrients. The eukaryotic flagellum is an ideal model for studying size sensing and control because its linear geometry makes it essentially one-dimensional, greatly simplifying mathematical modeling. The assembly of flagella is regulated by intraflagellar transport (IFT), in which kinesin motors carry cargo adaptors for flagellar proteins along the flagellum and then deposit them at the tip, lengthening the flagellum. The rate at which IFT motors are recruited to begin transport into the flagellum is anticorrelated with the flagellar length, implying some kind of communication between the base and the tip and possibly indicating that cells contain some mechanism for measuring flagellar length. Although it is possible to imagine many complex scenarios in which additional signaling molecules sense length and carry feedback signals to the cell body to control IFT, might the already-known components of the IFT system be sufficient to allow length dependence of IFT? Here, we investigate a model in which the anterograde kinesin motors unbind after cargo delivery, diffuse back to the base, and are subsequently reused to power entry of new IFT trains into the flagellum. By modeling such a system at three different levels of abstraction we are able to show that the diffusion time of the motors can in principle be sufficient to serve as a proxy for length measurement. In all three implementations, we found that the diffusion model can not only achieve a stable steady-state length without the addition of any other signaling molecules or pathways, but also is able to produce the anticorrelation between length and IFT recruitment rate that has been observed in quantitative imaging studies.

## INTRODUCTION

How does the cell know how big to make its organelles? This question has been puzzling cell biologists for decades. Cells must have a robust and efficient procedure for building organelles with a specific size and shape. The stochastic kinetics of polymerization typically leads to formation of structures with widely varying sizes in the absence of any size-dependent assembly or disassembly processes (1). But organelles are thousands of times bigger than the materials used to measure and build them. How can molecular pathways of assembly sense and respond to organelle size to yield organelles of a necessary size for proper function? This problem is extremely difficult to solve in the general case considering the many different types of organelles and their often highly complex structures. In order to simply the problem, we will just consider the eukaryotic flagellum. Flagella (also known as cilia) are long whip-like appendages protruding from certain cells, and are used for both locomotion and sensing. Unlike a prokaryotic flagellum, which is made of a tube of a single polymer, the eukaryotic flagellum is a more complex structure made of nine microtubule doublets underlying a structure of the plasma membrane. These doublets are nucleated by the basal body. The flagellum is the perfect organelle to model mathematically because it has a linear geometry: when it grows, it gets longer but not wider, making it essentially a one-dimensional organelle.

Here, we will consider the flagella of *Chlamydomonas reinhardtii,* a eukaryotic alga that has two flagella. When *Chlamydomonas* develop, their flagella grow with decelerating kinetics, ultimately leveling out to a steady-state length (2). This slow-down in growth suggests that some part of the flagellum-building mechanism can recognize when the flagellum is long enough. The present study examines how this might happen.

Most of the flagellum-building machinery is understood. To build a flagellum, cells use a process called intraflagellar transport, or IFT (3, 4, 5, 6). IFT, diagrammed in Figure 1A, is mediated by complexes of approximately 20 polypeptides called IFT proteins, which contain numerous protein-protein interaction domains capable of binding the building blocks of flagella such as tubulin and axonemal dynein arms. These IFT protein complexes associate into linear arrays known as “trains” (7,8). IFT trains are pulled to the distal tip by heterotrimeric kinesin-2 motors (9,10). Upon reaching the tip, the contents of the cargo add to the length of the flagellum. Flagella are thus undergoing continuous incorporation of new tubulin and other building blocks. To counter this, tubulin is continually removed from the flagellar tip at a constant, length-independent rate. Since this decay rate is constant, in order to achieve a steady state, the rate of IFT must be length-dependent (11,12).

**Figure 1.**
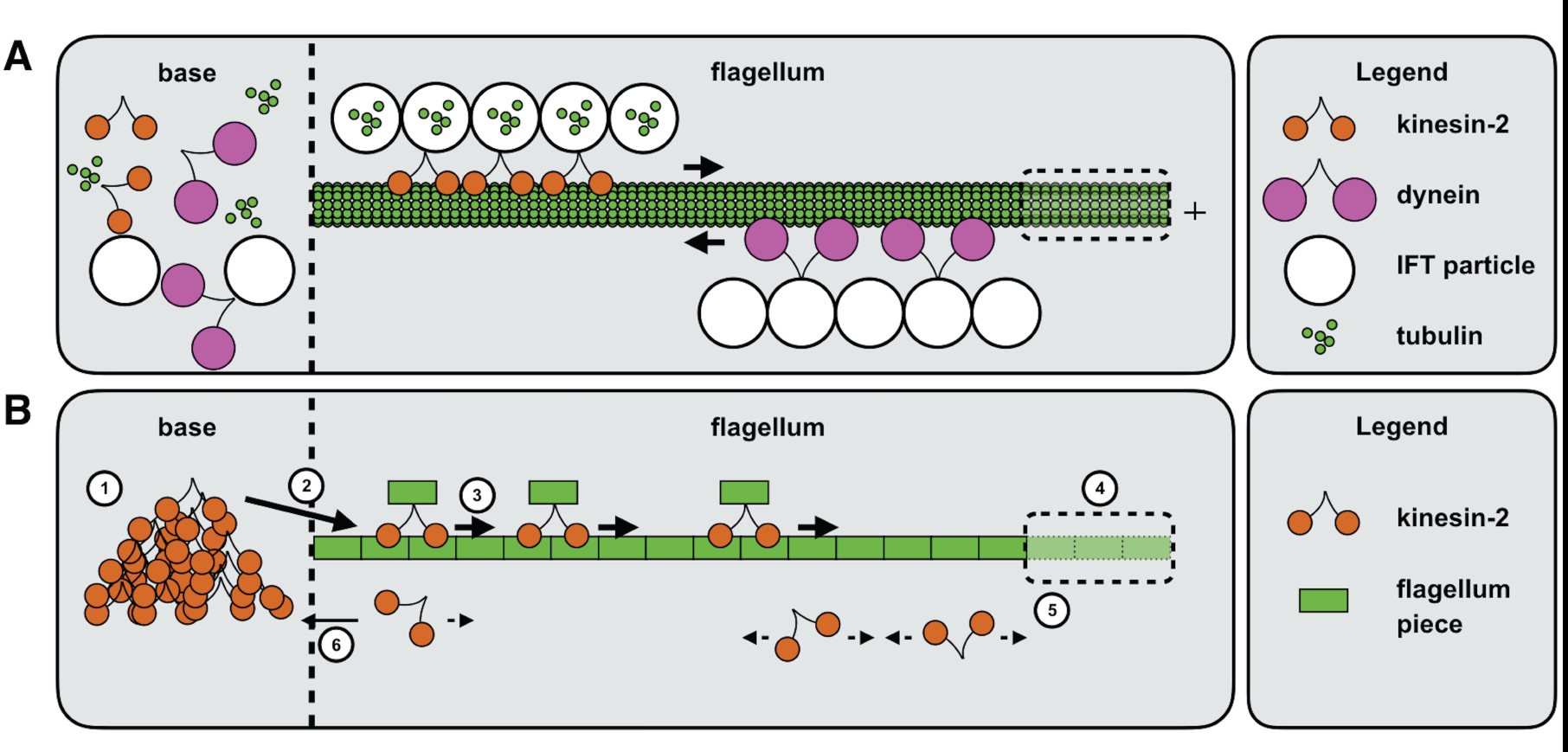
Agent-based model of IFT. (A) Diagram of IFT. Kinesin-2 motors form trains that carry IFT particles containing tubulin to the plus end of the microtubule bundle, the tip of the flagellum. Dynein motors carry the IFT particles back to the base. (B) Model version of IFT. Kinesin motors pile up at the base (1), and once the pile is large enough, some are injected into the flagellum with cargo (2). Each motor constantly moves towards the tip of the flagellum (3). Once they reach the end, they flagellum gets longer (4), and the kinesin motors unbind and diffuse (5). Once they diffuse back to the base, they are absorbed and re-enter the pile in the base (6). While this is happening, the flagellum is shrinking at a length-independent rate.

IFT trains are recruited from docking sites on the basal bodies (13) into the flagellum to begin transport through a process called injection. The physical mechanism of injection is unknown, but it is thought to involve IFT trains moving through some sort of selective pore or barrier similar to a nuclear pore (14, 15). While the molecular details of the injection process remain unclear, quantitative imaging studies (16) have revealed that motors are recruited into the flagellum according to a pattern of dynamics similar to how sand dropped onto a sandpile will fall off (avalanche) if the pile is high enough. For example, the more time elapses before a train is injected, the larger the train is, and the larger a train is injected, the more time will elapse before the next injection event. The sizes of the injection events are power-law distributed, similar to the size of avalanching events in sandpiles and other avalanching systems. These similarities suggest a simple model in which IFT proteins and motors accumulate at the basal body, gradually exerting more force on the pore until eventually a cluster of motors pushes through the pore, injecting a train (16). In such a scenario the rate at which motors accumulate at the base would ultimately be what determines the rate of injection.

Quantitative live cell imaging (16, 17) has shown that the rate of recruitment of motors is anticorrelated to the length of the flagellum. Furthermore, quantitative analysis of IFT cargo loading suggests that cargo loading is also length-dependent (18). These length-dependencies imply some kind of communication between the base and the tip. Perhaps some sort of additional signaling pathways have evolved that can sense length, transduce length into some form of molecular signal, and then use this signal to modulate the injection of IFT proteins at the base of the flagellum. Several possible models for length-sensing pathways have been described and analyzed (16, 19). Each of these models invokes additional molecular pathways that could transduce length into a signal that would gate entry of IFT particles through a pore. Is it possible, however, that no such additional pathway exists, and that the IFT machinery itself might be capable of responding to changes in flagellar length?

Here we consider a model that takes into account the return of motors from the flagella tip. IFT is a cyclical process: IFT trains and motors move to the tip, deliver cargo, return to the cell body, and then are re-injected (20). Experimental data has addressed how motors are recruited onto the flagellum, how motors get to the tip, and how the flagellum grows and shrinks. Two aspects of the IFT system that have been less intensively studied are how motors are sent to the pool at the basal body and what happens to the anterograde kinesin motors after they deliver their cargo to the tip. We propose a simple model to answer both of these questions: after dropping off their cargo, the motors unbind and diffuse back to the base, where they are then added back into the pool of accumulated motors waiting to be injected. The initial evidence for a diffusive return of the kinesin motor is the failure to observe processive retrograde traces in kymographs of IFT using GFP-tagged kinesin subunits (17), and the fact that when retrograde IFT is inhibited, flagella accumulate IFT proteins at the tip but not the kinesin motor (21). Direct tracking of individual trains by a novel bleach-gate method has shown that kinesin undergoes diffusion after dissociation from trains at the distal tip (22). In considering simple models for IFT that incorporate diffusive return of kinesin, we observed that the rate of diffusive return of kinesin motors to the pool at the flagellar base can serve as a proxy for flagellar length measurement, leading us to propose that the diffusion of the IFT kinesin motor may, itself, be the long-sought length sensor that regulates IFT injection.

In this paper, we investigate this hypothesis using models constructed at three different levels of abstraction: a fine-grained agent-based model that is analyzed using computer simulations, a stochastic process model that is investigated using linear algebra, and a coarse grained differential equation model that can be solved analytically. In the agent-based model, we explicitly model the flagellum and motors and run time dynamics simulations. In the stochastic process model, we construct a transition matrix and use its mathematical properties to determine a steady state. In the differential equations model, we solve the steady state form of the diffusion equation with boundary conditions that incorporate active delivery of IFT to the tip and diffusive return to the base. Each model is detailed below.

## AGENT-BASED MODEL

As a starting point to look for potential length dependencies in the IFT system, we implemented a simplified model of the individual components of the system (Figure 1B) and asked what predictions this model might make about length dependence. We built an agent-based model to simulate kinesin and microtubule growth dynamics through stochastic rules grounded in biochemistry. Specifically, we used Python’s built-in object oriented programming methods to explicitly model individual motors and the flagellum they populate.

The flagellum has attributes including length and environmental variables including decay rate and diffusion coefficient. The motors each have attributes including position, transport speed, if they are bound, and if they are on the flagellum or in the base. To simulate dynamics, we cycle through each motor and test a series of conditionals to determine how it should adjust its position. If it is on the flagellum and bound, it moves a constant rate forward. If it reaches the tip of the flagellum, it unbinds, and the flagellum grows by the designated growth increment. If it is in the flagellum and unbound, it moves randomly to the left or to the right. If it is unbound and reaches the base, it is absorbed into the base and becomes inactive. At each time step, we count the number of motors in the base, and if that value is greater than a variable for avalanche threshold, we use a Weibull distribution to determine how many should avalanche out and move into the flagellum, and reactivate into active transport. We chose a Weibull distribution because it can fit the long-tailed distribution of train sizes that have been experimentally determined (16). The Weibull distribution has a multiplicative constant that we set to the difference between the number of motors in the base and the threshold for avalanching, plus a constant we could vary. Meanwhile, at each time step, the flagellum shrinks by the decay rate constant. Table 1 lists parameters we used, and how we obtained the values used for simulation.

**Table 1.**

| Parameter | Default value | How value was obtained | Notes |
| --- | --- | --- | --- |
| Number of motors | 200 | Marshall et al, 2001 (11) |  |
| Active transport speed | 2 $\mu\text{m/s}$ | Chien et al., 2017 (22) | |
| Growth size per motor | 1.25 nm | Marshall et al., 2001 (11) |  |
| Decay rate | 0.01 $\mu\text{m/s}$ | Marshall et al., 2001 (11) | |
| Diffusion coefficient | 1.75 $\mu\text{m}^2/\text{s}$ | Chien et al., 2017 (22) | |
| Weibull distribution<br>power | 2.85 | Ludington et al., 2013 (16) |  |
| Weibull distribution<br>constant | 10 | Arbitrary |  |
| Avalanche threshold | 30 motors | Ludington et al., 2013 (16) |  |
| Binding on | 0 | Arbitrary | Probability for<br>each diffusing<br>motors to bind to<br>the flagellum |
| Binding off | 0 | Arbitrary | Probability for<br>each bound motor<br>to unbind from the<br>flagellum |

This model lets us consider the journey of a single motor (Fig. 2A). In this example, it starts at position 0, with the bound parameter set to True. The conditional that checks if it is bound commands its position to increase by the active transport step size. This process continues until the position of the motor is equal to the length of the flagellum. This position represents the tip, and at this stage, the motor’s bound parameter is changed to False, and the length of the flagellum is increased by the build size parameter. In the next time step, the conditional that checks if the motor is bound sees that it is not bound, and this time it adjusts its position by the diffusion length multiplied by either 1 or −1, determined randomly. This simulates the randomness of diffusion. Once its position reaches 0 (the base), its Boolean value stating whether it is active (meaning, on the flagellum or diffusion) is set to False to indicate absorption to the basal pool. Every time step, a random power law number generator determines how many motors that are inactive at the base are injected onto the flagellum. This process then repeats for the remainder of the simulation. By saving the flagellum’s length after each time iteration, we can plot its length over time curve shown in figure 2B.

**Figure 2.**
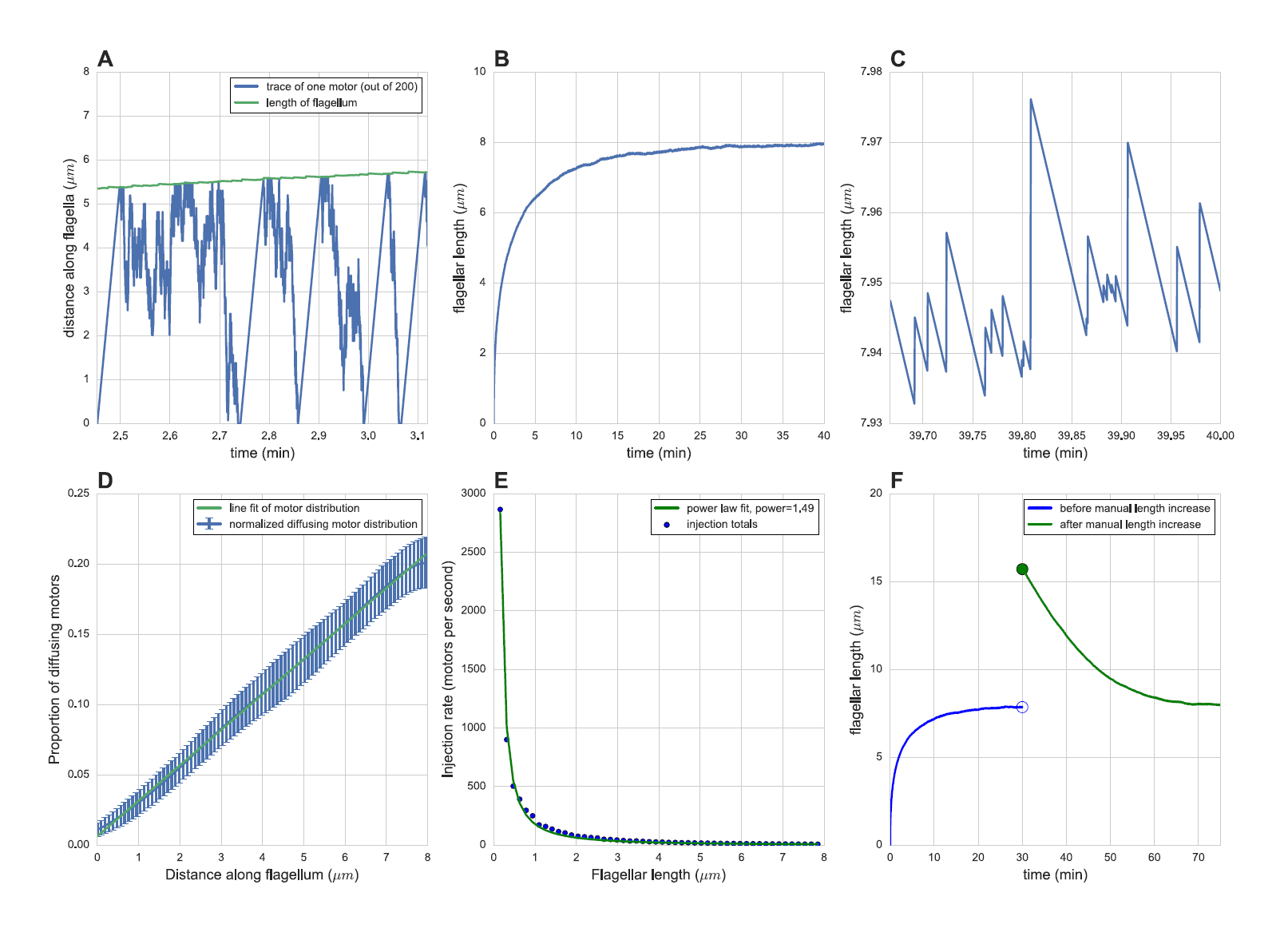
Results of agent-based simulation. (A) (Blue) journey of a single motor in a zoomed-in window of the flagellum’s early growth, (green) flagellar length. (B) Length over time in simulated minutes. (C) Zoomed in window of the flagellum’s length over time curve in the steady state regime. (D) (Blue) Distribution of diffusing motors along flagellum using the average of 103 simulations with identical parameters, then applying a Gaussian kernel density function to the means, (green) linear fit. (E) Plot of injection size as a function of flagellar length. The points were generated by simulating 10 cells, taking their injection times and sizes, and binning them into measurements of average injection size per unit time in each of the 50 evenly-spaced bins. (F) Stability of length control system. Plot shows simulation in which length was manually increased to double its steady state length at t=30 min. (Blue) is before the manual increase, (green) is after, showing restoration to initial steady state length. The time step in each simulation was 0.01 seconds.

Simulations over time show that this system allows the flagellum to grow to a defined length with decelerating kinetics (Fig. 2B). This diffusion-based control scheme is robust and works for a wide range of parameters.

Because motors undergo random motion as they return, and are released from the base in a way that depends on the time history of their return, it is expected that flagellar growth rates will fluctuate, and indeed our simulations confirm that the length does indeed fluctuate around a steady state average length (Fig. 2C). By counting motors in different states, we can ask how the pool of diffusing motors is distributed along the length. We find that the probability of finding a motor at a give distance from the tip is approximately linear, consistent with the expected form of a diffusional gradient at steady state (Fig. 2D).

Having found that the simple agent-based model of diffusive kinesin return is able to produce a defined flagellar length, the key question is whether the length-dependence of IFT injection can be recapitulated. As shown in Figure 2E, the average injection size per unit time of injected IFT trains in the simulation shows an inverse dependence on flagellar length, as previously reported in experimental measurements (16, 17).

The length control system modeled here is stable, as indicated by simulated experiments in which the length is transiently perturbed. As illustrated in Figure 2F, transient elongation of the flagellum is followed by a shortening back to the steady state length. Once the flagellum reached steady state, we manually doubled its length and resumed the simulation until the flagellum reached steady state again. This implies that the steady state length is determined by the input parameters rather than the transient state of the flagellum.

## TRANSITION MATRIX MODEL

In order to understand why this diffusion-based mechanism actually works and how it depends on parameters, one approach would be to explore the entire parameter space of the model using exhaustive methods, but this would require a prohibitive number of simulations. We therefore seek a more abstract model that can be analyzed mathematically to yield a more intuitive understanding of why the model works the way it does. To this end, we modeled the flagellum as a column vector N(t), with each element in the vector representing the number of motors at that location processing along the flagellum at time t. We then extended that vector to twice the length of the flagellum, with each element in the second half representing the number of motors diffusing at the corresponding location. Finally, we extended the vector by one element to represent the number of motors in the base. We can then represent the dynamics of the entire system using a stochastic matrix M such that M*N(t) = N(t+1).

Figure 3A shows an example transition matrix M representing the dynamics of a flagellum of length 4. To construct M, we need to consider several constraints. First, the number of motors in the system must be conserved, so the sum of the elements in the state vector N(t) must remain constant throughout all t. The columns can be thought of as the spread of a point source after one time step. Specifically, if the value of the state vector component at position j at time t is n_j_, the transition matrix will redistribute those n_j_ motors into a new distribution, governed by the values in M. Since every motor needs to end up in some position (given conservation of total motor number), the entries in the whole column must sum to 1. The condition that each column in M must sum to 1 defines M as a left stochastic matrix. This property of the matrix will help us later determine the steady state of the system and solve the length control problem.

**Figure 3.**
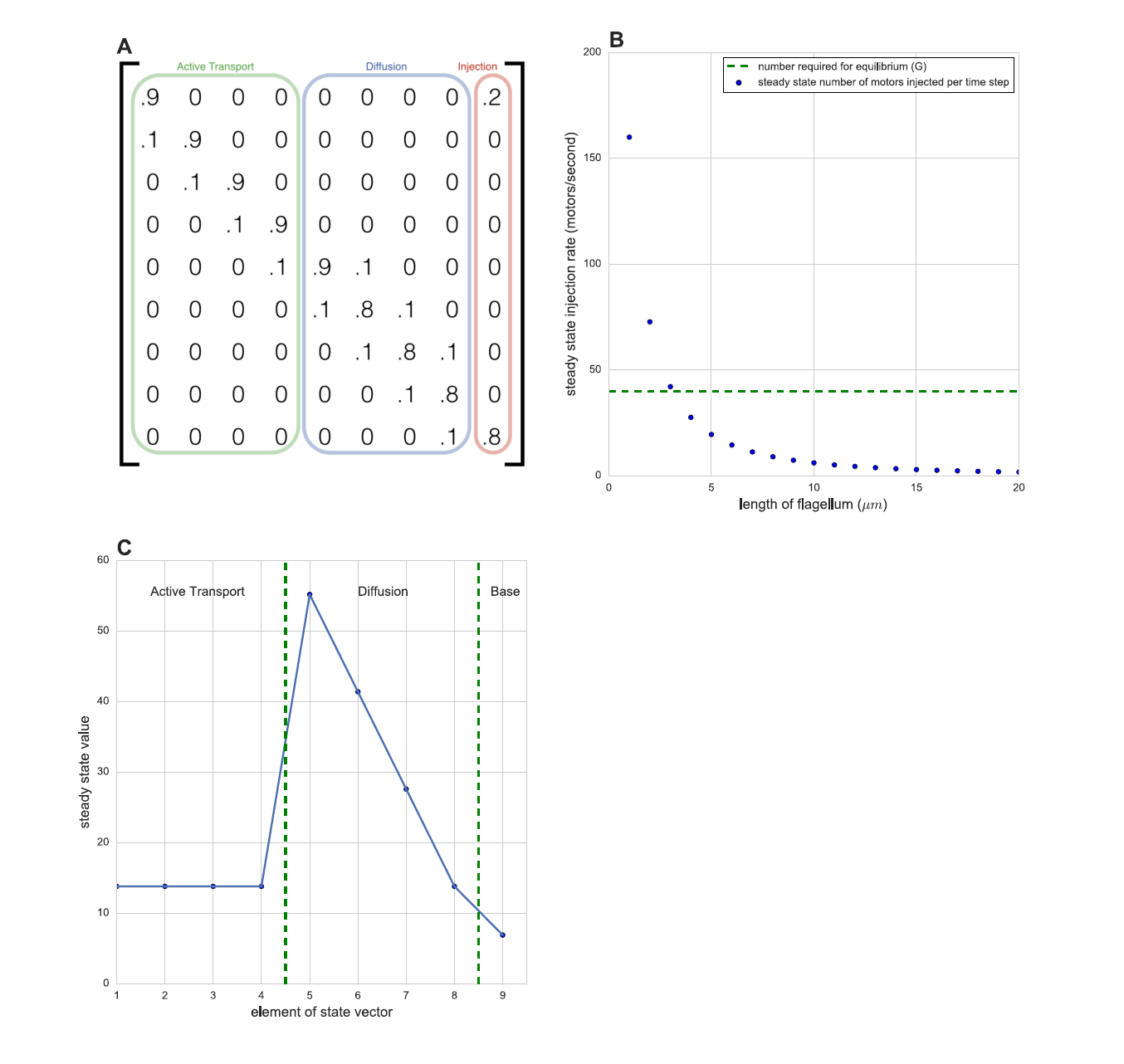
Markov matrix model. (A) Example of a transition matrix, here with length 4, active transport rate of 0.1, diffusion spread of 0.1, and injection rate of 0.2. The relative sizes of the active transport rate and diffusion rate are roughly equal to the biological parameters used in the agent-based model, but the injection rate is simplified to a length-independent proportion. Based on the active transport and diffusion parameters, this matrix advances a state vector forward in time by 0.05 seconds. (B) Steady-state injection rate as a function of length compared to the value G required for equilibrium. (C) Steady state number density (principal eigenvector) for one set of parameters. x = 1:4 is active transport, x = 5:8 is diffusion, x = 9 is base. Note that the eigenvector can be scaled to an arbitrary magnitude, here it makes sense to normalize it to sum to the number of motors in the system, which we set to 200 for consistency with the agent-based model.

Second, the matrix must simulate active transport for the top half of the state vector, diffusion for the bottom half, and absorption/recruitment to send motors from the bottom value to the top value. Since we constructed the state vector such that the first L values represent bound (i.e. transporting) motors, the top left quadrant of the transition matrix M will represent the active transport dynamics. Active transport is simply moving some percent of motors one unit forward and keeping the remaining motors at their current position at each time step, so the active transport quadrant of the matrix will have positive values on the diagonal and one position under the diagonal.

The diffusion region of the transition matrix must apply to motors that have moved past position L in the state vector. This means that the lower right quadrant of the transition matrix M must simulate the dynamics of diffusion. We can incorporate the random walk nature of diffusion into this matrix by stating that the probability of staying in the same position is high, and the position of moving one position to either side is low. This simulates the Gaussian spread of a diffusing point source after a small time (we keep the time small so there is a negligible chance of diffusion two units away).

Notice that the first column incorporates the reflecting boundary condition that motors cannot go past the tip, so the odds of staying at the tip are the odds of not moving anywhere (here 0.98) plus the odds of moving past the tip and bouncing off (here 0.01). Also note that the way our state vector is constructed, motors diffusing in the direction of the base are going down the state vector towards lower rows. This matches the order in which vector elements representing diffusing kinesins are specific in the state vector

With the aforementioned elements of M specified, we are able to represent how the motors can actively transport to the tip, unbind, diffuse back to the base, and absorb at the base so that motors enter the inactive pool. We still need to add the final element of our dynamics into the matrix: injection. A simple way to do this is to assume that at each time step, the base sends p percent of the motors in the base back to the flagellum for active transport. This means that 1-p represents the proportion of motors that stay in the base. Such an assumption is a simplified representation of the quasi-periodic avalanching process, and may need to be relaxed in future simulations. The last column in M represents the spread of motors that were previously at the base. To incorporate avalanching and recruitment into this column, we simply make the column [p 0 0 … 0 0 1-p]^T^, where p is the probability of a motor being injected.

Now all the columns in the matrix sum to 1, so the condition for being a stochastic matrix are satisfied. The probability of different states evolves in a strictly deterministic manner determined by successive matrix multiplications. For example, if the diffusion half of the state vector is [0 1 0 0]^T^, applying M will result in a new state vector whose elements are real numbers in the range 0 to 1 that represent the probability of a motor occupying that position in the state vector. This makes sense physically in the assumption that there are a large number of motors in the system, and since the number is on the order of 200 motors, this is a reasonable approximation.

One limitation of this construction of the transition matrix is that it assumes a constant flagellum length. The length determines the size of the matrix, so to simulate length dynamics over time, we would need to continuously alter the size of the matrix. To avoid this inconvenience, we can instead directly calculate the steady state behavior as a function of flagellar length. The steady state solution *N*_*SS*_ must satisfy *M * N*_*SS*_ = *N*_*SS*_, so *N*_*SS*_ is an eigenvector of *M* with eigenvalue 1. The Perron-Frobenius theorem states that the largest magnitude eigenvalue of stochastic, nonnegative, and irreducible matrix is always simple and equal to 1. Our motor transition matrix is stochastic (i.e. Markov) because the columns each sum to 1. It is nonnegative because all values are greater than or equal to zero. Finally, it is irreducible because each node has a path to get to every other node after some number of time steps. For example, a motor in the middle of active transport has a path leading through every subsequent active transport node, then it connects to a diffusion node, and each diffusion node is connected to a subsequent diffusion node, the last one connects to the base node, which connects to the first active transport node. This means we can apply the Perron-Frobenius theorem for nonnegative irreducible matrices to this stochastic matrix, proving that the eigenvalue of 1 always exists and is unique, and corresponds to a principal eigenvector corresponding to the steady state number distribution (N_SS_ in our example). This also means that the system is robust, and all sizes of the matrix M will yield a steady state solution. Because all other eigenvalues must have magnitudes less than 1, the corresponding eigenvectors will decay in any superposition state, so the same steady state solution will always be attained regardless of initial state. No change to the numerical values of the parameters in the model will cause the matrix M to violate the conditions of the Perron-Frobenius theorem, hence there will always be a unique steady state no matter how the parameters are altered. This property of stable length control is a robust feature of the system.

This method represents IFT in a flagellum at any fixed length, which determines the size of the state vector and transition matrix. The flagellum grows when motors with cargo reach the tip, and shrinks through a constant, length-independent decay. When the number of motors arriving at the tip times the growth per motor equals the decay in some time interval, the net length change will be zero. Since motors in active transport move at a constant rate, the number of motors injected into active transport is the only factor that controls the number arriving at the tip per second. This value can be expressed as the number of motors in the base multiplied by *p,* the fraction of motors in the base that get injected into active transport. We can therefore define the critical rate of motors that must arrive at the tip to maintain a steady state length as *G* = *d*/(δ*L * p*), where *d* is the decay rate and δ*L* is the growth increment when a single motor reaches the tip. The value of the steady state number density vector *N*_*SS*_ in position (2L+1) is the number of motors at the base. This means that when N_SS_(2L+1) > G, there are enough motors at the tip that the flagellum will grow. If N_SS_(2L+1) < G, there are too few motors to counteract the decay, so the flagellum will shrink. This means that when N_SS_(2L+1) = G, the growth factor from motors at the tip perfectly cancels the decay rate. Therefore, when N_SS_(2L+1) = G, the matrix is the right size to encode a flagellum that reaches steady state length.

We can find this matrix by creating transition matrices corresponding to a range of lengths, finding each matrix’s principle eigenvalue, and examining the value of the corresponding eigenvector at position (2L+1). Figure 3B shows the values at this position as a function of L. The horizontal line represents the value of G given by the default parameters in the agent-based model. The matrix that intersects the line at G is the one with the steady state length. The difference between this steady state length and the result from the agent-based model may be explained by the different implementation of avalanching between the models. Note the inverse relationship between injection rate and flagellar length, matching experimental results (16). A possible future direction for this model is making the separation between elements in the matrix correspond to a smaller unit of length, or perhaps a continuous differential equation, allowing us to precisely predict final length. The equilibrium here is stable, reiterating the point that the length would modulate until it reaches steady state. It also means that this system is robust, because any parameter adjustment would retain the stable equilibrium. This model also predicts that the gradient of diffusing motors is linear (Fig. 3C), like in the agent-based model. The benefit of the matrix model in addition to the agent-based model is that it provides an intermediate level of scale that proves stability and robustness, and that it is efficient to vary biochemical parameters and find the steady state solution.

## DIFFERENTIAL EQUATIONS MODEL

The stochastic process model described above provides a simplification of the initial agent-based model, but it still requires numerical solutions to find steady state distributions of motors. We therefore investigate an even more idealized model that will allow us to solve for the steady state solution analytically, so as to determine the influence of key parameters on system behavior. If we make the assumption that active transport time and expected time delay of injection is small relative to the timescale of diffusive return, we can model this system as a diffusion problem with a constant source of free motor protein at the tip of the flagellum and a sink at the base. If we also assume that no diffusing motors re-bind to the flagellum, we can apply Fick’s first law of diffusive flux in steady state. This law strictly applies to steady state, however we can still use it to study the dynamics of flagellar growth by invoking a separation of timescales. We assume that the timescale of flagellar length changes due to growth and shrinkage, which happens on the timescale of minutes to hours, is slow relative to the timescale over which diffusion establishes a stable gradient, such that the system can be viewed as being in a quasi-steady state. (This similar to the classic statistical mechanics problem of slowly expanding a box containing gas: when the expansion of the box is slow, the system is reversible and equilibrium statistical mechanics theory can be applied. A simple validation of this is that a single motor reaching the tip increases the length by 1.25nm in our simulation, and it takes 4.5e-7 seconds for a diffusing motor’s mean square displacement to equal 1.25nm, which is negligible compared to the time it takes to diffuse back to the base, roughly 18 seconds).

The strategy for deriving an expression for steady state length is to determine the expected flux of diffusing motors arriving at the base, equate the flux to the number of motors diffusing from the tip (following our assumptions that injection time and active transport time are very small compared to diffusion time), convert that flux into a dynamic growth term, and then find the steady state at which this growth is balanced with the decay term.

The resulting expression for steady state length is the following:

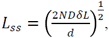

Equation 1,

where *N* is the number of diffusing motors, *D* is the diffusion coefficient, δ*L* is the increment of flagellar growth when a motor reaches the tip, and *d* is the decay rate.

It can be shown from first principle random walk distance distributions that the time it takes to move a root-mean-square distance L is:

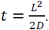

The current of motors *I* reaching the base is equal to the number of diffusing motors *N* divided by the average time it takes to diffuse to the base

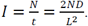

In the approximation in which motors that have reached the base immediately transport back to the tip, the flagellum grows by the current of motors reaching the base multiplied by the growth increment per motor δ*L*. The competing decay term *d* is length-independent.

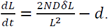

At steady state 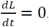, so it is simple to solve for the steady state length *L*_*SS*_.

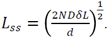

An identical result can be obtained by solving the diffusion equation for appropriate boundary conditions and then expressing the motor return rate in terms of the flux at steady state.

This predicts that the steady state length of the flagellum is proportional to the square root of its diffusion coefficient, motor number, and unit length increase per motor. It also predicts that it is inversely proportional to the square root of the decay rate. Note that since the model proposed does not invoke any unknown transducer molecules or pathways, but instead directly represents all of the molecular players, there is no need for any undetermined constant of proportionality.

Note that *N* here represents number of diffusing motors, not total motors. In our assumption that injection frequency and active transport are fast, *N* is equal to the number of diffusing motors, and when these assumptions break, there should be some correction term, perhaps *N*_effective_=*N*_*total*_ – (threshold for avalanching). Unless otherwise specified, in our simulations we used threshold=1, so *N*=199 out of 200 motors.

By running simulations in the agent-based model over a range of parameters, we can verify that this relation matches the results of fine-grained agent based simulations. (Fig 4). To simulate our assumptions, these simulations have an avalanching threshold of 1 and an active transport speed of 200 μm/s (enough to go the entire length of the flagellum in one time step). This deals with the regime of high active transport velocities, which is neglected by the Markov matrix model. To correct equation 1 in the future to include low velocities, we would need another small correction to *N*, because slow walkers are essentially motors in the system that are not diffusing. The similarity between the curve fits and the simulated lengths indicate that equation 1 accurately describes the length of diffusion-regulated flagella.

**Figure 4.**
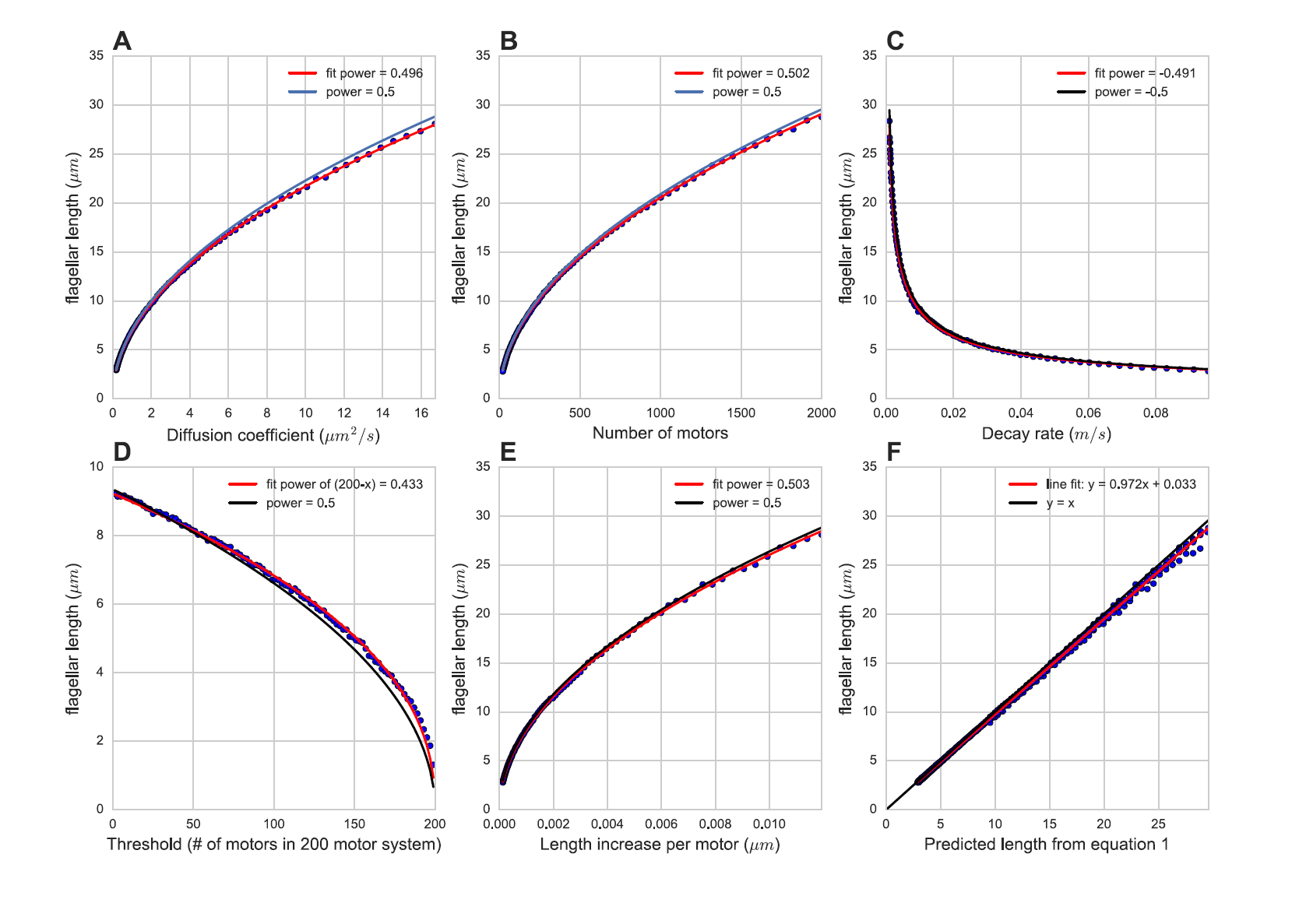
Comparison of analytical solution of diffusion equation to agent-based model. Each plot shows the lengths given by equation 1 and agent-based simulations by varying a single parameter at a time. The varied parameters are: (A) diffusion coefficient *D*, (B) number of motors *N*, (C) decay rate *d*, (D) avalanching threshold, (E): length increase per motor δ*L*, (F): all parameters, using the data from panels A, B, C, and E, and multiplying the variables to match equation 1, then comparing to final length simulated by the agent-based model. The red curve in each is the best-fit curve to the curve (a*x)^b^ (except the threshold graph, which is (a*(200-x))^b^, and the value for the fit power *b* is displayed in each legend. The blue curve is the predicted curve given by equation 1. The points for panels (A), (B), (C), and (E) were uniformly sampled in log space, so there are the same number of points between the default and one order of magnitude below as there are between the default and one order of magnitude above.

## DISCUSSION

### Diffusion as a ruler

In this model of length sensing, the cell is not sensing length directly, but it is converting a biochemical signal that obeys the laws of diffusion and using it as a proxy for length measurement. This is similar to a chemical reaction in which a chemical X has an assembly term and a degradation term. The concentration of X over time is given by a simple differential equation, and the steady state concentration is determined by a combination of biochemical parameters. The flagellum is a similar system because the length has assembly and disassembly terms, and here we predict which specific biochemical parameters are involved (equation 1). There is a competition between a growth flux term (δ*L * N * D*) and a decay term *d*. It is important to note that the square root in equation 1 comes from the geometry of the system.

### Relating model to genetics of length control

The simple mechanism modeled here is sufficient to explain length-dependent IFT injection and stable length control without needing to invoke any new molecular players beyond those already known. But this does not mean that the model works independently of molecular entities. All of the model parameters are determined by the biophysical and enzymatic properties of the known molecular component of the IFT system. It is to be expected that mutations in these molecules can alter flagellar length in predictable ways, potentially allowing the model to help interpret the mechanistic basis of previously described flagellar length-altering mutants.

The diffusion constant of kinesin is mainly a property of the size of the molecule and the viscosity of the flagellar matrix, and is thus unlikely to be dramatically altered with point mutations. But it is not hard to imagine that mutations might alter the dynamics of the injection system at the base. Previous research shows that the *lf4* mutant makes the flagellum longer and increases the injection rate but without eliminating the length dependence of injection (16). Such a phenotype could correspond to lowering the threshold of motor buildup required for injection avalanching, which is a parameter in the agent-based model. High thresholds lead to lower injection frequency and lower steady state length, and low thresholds lead to higher injection frequency and higher steady state length. This breaks the assumption of equation 1 that injection is instantaneous, and essentially it lowers *N* by reducing the fraction of motors in diffusion. This implies that it is possible that the LF4 gene controls the threshold for how big the pile can be before an avalanche occurs.

Another mutant that we can examine is the FLA10 gene, which codes for the kinesin motors (9). Temperature-sensitive *fla10* mutants with intact flagella start to lose their flagella when the temperature shifts into the region that disables FLA10 (9). Growth of *fla10* mutants at intermediate temperatures, which partially disable the motors, leads to intermediate steady-state flagellar lengths (11). In our model, this translates to a reduction in N, the number of motors in the system. We note that the square-root dependence of steady state length on motor number (equation 1) means that length will decrease sub-linearly with decreasing motor number. To reduce length by a factor of 10 would require a reduction in motor number by a factor of 100. Since motors reaching the tip and delivering cargo is the only mechanism in the model for flagellum growth, removing every motor makes the flagellum shrink to zero. This is another prediction of equation 1.

### Comparison with other studies

A recent study on mouse axons (23) studies the diffusion of kinesin motors as a mechanism for recycling. Their model for simple diffusion has the same linear distribution of diffusing motors, but they find that the diffusing motors have a nonzero binding rate onto the flagellum from diffusion, and therefore the number distribution is exponential. The mouse axon system has a fixed length, but their work provides an example in biology of diffusion and recycling of kinesin.

Models based on diffusion as a length measurement system have been proposed by Levy (24) and by Ludington (16). In the model by Levy, the proposed source of the diffusing molecule was the base, not the tip, and it was assumed that the diffusing species directly affected assembly, as opposed to our model in which the diffusing molecule affects transport. In the Ludington 2013 model, RanGTP was the diffusing substance, and the link to injection was indirect, requiring a gating of entry by activated Ran. In the diffusion model investigated in Ludington 2015, the identity of the diffusing molecule was not specified and again a transducer system was assumed to couple the diffusive molecule to the injection system (19). Finally, we note that while a strength of our model is that length can be sensed and converted into length-dependent IFT injection without the need to invoke any other molecular players, it has been shown that kinases inside the flagellar compartment do show length-dependent activity (24, 25). Likewise, flagellar disassembly can become length dependent when flagella grow outside of a normal length range (27). It is interesting to consider whether these molecular activities may be dependent on IFT injection or diffusive return.

### Future Prospects

A fundamental puzzle of flagellar length control has always been how the organelle can measure length. Our prior results indicated that IFT injection was length dependent but did not explain the origin of the length dependence, thus raising the possibility that some complex length-measuring molecular pathway may exist. The results presented above establish that diffusive return of kinesin motors is, at least in principle, capable of providing a length measurement system for regulating IFT injection as a function of flagellar length, without requiring any additional regulatory or sensing components. In other words, the IFT system may contain its own measurement method based on the physics of diffusion. It is interesting to consider whether this type of measuring system could be at work in other linear cellular structures such as microvilli or microtubules.

## AUTHOR CONTRIBUTIONS

N.H. wrote the simulations. N.H., M.T., and W.M. developed ideas and worked out math. N.H. and W.M. wrote the manuscript.

## ACKNOWLEDGEMENTS

We would like to thank Greyson Lewis for help with the math and derivations and Ahmet Yildiz for sharing results ahead of publication. This work was supported by NIH grant GM097017.

## CODE AVAILABILITY

Code for agent-based simulations and Markov matrix simulations is available at https://github.com/nathendel/Hendel-et-al-2017.

